# NAc-DBS corrects depression-like behaviors in CUMS mouse model via disinhibition of DA neurons in the VTA

**DOI:** 10.1101/2021.12.01.470503

**Authors:** Song Nan, Gao Yan, Lu Shanshan, Yang Shenglian, Yuan Chao, Sun Wenyu

## Abstract

Major depressive disorder (MDD) is characterized by diverse debilitating symptoms that include loss of motivation and anhedonia. If multiple medications, psychotherapy, and electroconvulsive therapy fail in some patients with MDD, their condition is then termed treatment – resistant depression (TRD). MDD can be associated with abnormalities in the reward–system–dopaminergic mesolimbic pathway, in which the nucleus accumbens (NAc) and ventral tegmental area (VTA) play major roles. Deep brain stimulation (DBS) applied to the NAc alleviates the depressive symptoms of MDD. However, the mechanism underlying the effects of this DBS has remained elusive. In this study, using the chronic unpredictable mild stress (CUMS) mouse model, we investigated the behavioral and neurobiological effects of NAc – DBS on the multidimensional depression – like phenotypes induced by CUMS by integrating behavioral, *in vivo* microdialysis coupled with high-performance liquid chromatography – electrochemical detector (HPLC – ECD), calcium imaging, pharmacological, and genetic manipulation methods in freely moving mice. We found that long–term and repeated, but not single, NAc–DBS induced robust antidepressant responses in CUMS mice. Moreover, even a single trial NAc–DBS led to the elevation of the γ–aminobutyric acid (GABA) neurotransmitter, accompanied by the increase in dopamine (DA) neuron activity in the VTA. Both the inhibition of the GABA_A_ receptor activity and knockdown of the *GABA_A_*–*α1* gene in VTA–GABA neurons blocked the antidepressant effect of NAc–DBS in CUMS mice. Our results showed that NAc– DBS could disinhibit VTA–DA neurons by regulating the level of GABA and the activity of VTA–GABA in the VTA and could finally correct the depression–like behaviors in the CUMS mouse model.

## Introduction

Depressive disorders affect approximately 5% of the population in any given year [1]. A significant number of patients with major depressive disorder (MDD) remain inadequately treated and display nonresponse or partial response after all treatment options have been explored [2, 3]. This treatment-resistant depression (TRD) is particularly associated with a great economic burden [4], high social and familial impact [5], and personal suffering. New hope has now been given to these individuals by clinical studies demonstrating the long-term effects of high-frequency deep brain stimulation (DBS) in terms of improving the depressive symptoms of helplessness, anhedonia, and anxiety, thus enhancing quality of life [6–8].

Promising results have been obtained by DBS to the nucleus accumbens (NAc, NAc-DBS), which has induced solid antidepressant, anxiolytic, and antianhedonic effects in patients with TRD [7]. However, continuous intensive stimulation at a relatively high-frequency was needed [1]. The unique position of the NAc in the brain enables it to act as a gateway for the transmission of information from emotional centers to motor control regions [9]. Functionally, the NAc processes reward and pleasure information and is dysfunctional in patients suffering from depression [10], suggesting a potential role in the depressive symptoms of anhedonia [11–13]. This is clinically relevant as anhedonia is one of the core symptoms of depression and reflects a lack of reward and reward-motivated behavior [14].

The loss of motivation and anhedonia observed in MDD can be associated with abnormalities in the reward-system–dopaminergic mesolimbic pathway, in which the NAc and ventral tegmental area (VTA) play major roles [15]. Several lines of evidence have indicated the involvement of the dopaminergic system in the mechanisms underlying depression [16]. One of them demonstrated that patients with depression show a decrease in dopamine (DA) neurotransmission and a reduction in the cerebrospinal fluid levels of homovanillic acid, a DA metabolite [17, 18]. Additional support comes from the finding that there is an association between the alterations in DA levels achieved by pharmacological means and the level of depressive symptoms [17, 19]. A second source of evidence arises from the exploration of the pathways of the dopaminergic system. The NAc receives a dense dopaminergic input from midbrain areas, such as the VTA. In turn, the NAc provides one of the most prominent projections to the VTA [15, 20]. Findings from animal studies have indicated the NAc–VTA pathway as a mediator of reward-driven, pleasurable behavior [6, 21]. Detrimental changes in such behavior, which are exhibited by individuals with depression, further strengthen the case for the involvement of the dopaminergic system.

The mechanism of action of DBS has remained elusive. Early observations in patients with Parkinson’s disease led to the proposal of a functional blockage of the target region by DBS [22, 23] through the activation of γ-aminobutyric acid (GABA) related neurotransmission and increased release of the inhibitory neurotransmitter GABA, consequently modulating dopamine release [24, 25]. The NAc includes a cell cluster only for GABAergic neurons in the limbic system and is thought to participate in the brain reward circuit [26]. Additionally, inhibitory synapses comprise 50%–80% of all midbrain dopamine neuron synapses [27], which highlights the importance of GABAergic transmission in the VTA [20]. Thus, it is thought nowadays that GABA or GABA receptors are involved in the effectiveness of DBS and cause axonal activation and neuronal inhibition [6, 23, 28]. However, it remains unclear whether NAc stimulation operates via similar pathways [25]. Therefore, clinically relevant animal models are used to study the neurobiological mechanisms underlying the beneficial effects of NAc-DBS.

In this study, we used mice subjected to chronic unpredictable mild stress (CUMS), which is a condition that is used to induce behavioral alterations resembling MDD in humans [29], and provided the first comprehensive behavioral evaluation of NAc-DBS in the CUMS mice. We also aimed to explore the neurobiological mechanisms underlying NAc-DBS for intervention in the dopaminergic reward circuit and the attenuation of depressive symptoms.

## Materials and methods

All experiments involving animals were performed in accordance with protocols approved by the Institutional Animal Care and Use Committee of the Academy of Military Medical Sciences.

### CUMS model

We used a CUMS regimen that followed the procedure originally described by Willner et al. [30] and that was subsequently adapted to mice [29]. This stress model involves the repeated application of mild physical and psychological stressors. The CUMS paradigm provides a model of a chronic-depressive-like state that develops gradually over time in response to stress. Hence, CUMS is considered more naturalistic in its induction [31]. Here, we subjected the mice to different types of stressors several times a day for 4 weeks in a chronic, inevitable, and unpredictable way. For ethical reasons, the stress procedure did not involve food and water deprivation or immobilization. The stressors used here were as follows: damp sawdust; changing the sawdust; placement in an empty cage or an empty cage with water covering the floor; placement in a soiled cage with an aversive odor; cage tilting (45° for 12 h); noise stress (15 min); inversion of the 12:12 h light:dark cycle; lights on or off for a short time during the dark phase or light phase, respectively; foot shock (150 mA, 30 min); and ice-water swimming (15°C, 5 min). We administered these stressors in a pseudorandom manner so that they could occur at any time of the day or night, and we changed the stressor sequence weekly to ensure that the stress procedure was unpredictable. During the behavioral tests, the stress procedure was slightly modified: we reduced the number of stressors applied during the light period to avoid interference with the tests. In addition, we did not subject test mice to any stressors for 12 h before the behavioral tests. Nonstressed mice were left undisturbed in their home cages. In all experiments, the first 2 weeks of CUMS were DBS-free, whereas the second 2 weeks of CUMS also included the administration of DBS. To determine the behavioral effects of the CUMS regimen and DBS treatment and to confirm the induction of depression-like behaviors, we examined sucrose consumption, tail suspension, and forced swimming in mice. We also measured locomotor activity using an open field test. We isolated nonstressed animals for 1 day before the open field test to match the conditions used for the CUMS mice.

### Deep brain stimulation

Before surgery, 6-week-old C57BL/6J mice were handled for 7 days to habituate them to the experimenter and the stimulation procedure. Subsequently, we anesthetized the mice intraperitoneally (pentobarbital sodium, 0.7%, 10 μL/μg) and, using a stereotaxic frame, implanted electrodes (bipolar, two parallel tungsten wires, twisted, 0.22 mm in diameter) reaching the bilateral NAc (coordinates: 1.18 mm anterior, ±1.35 mm lateral, and −4.50 mm ventral to bregma). After 5–7 days of recovery and 2 weeks of a 4-week CUMS regime, we subjected the mice to high-frequency stimulations (100 Hz, 100 μA, 100 μs pulse width) daily for 1 h per day (30 min/unilateral) (i.e., NAc-DBS). Thus, stimulations took place from the second week of CUMS, and they continued for another 2 weeks (i.e., the end of CUMS). We controlled current intensity using a constant-current isolated stimulator, and we synchronized our stimulation protocols using Master-8 (A.M.P.I., Israel). We also placed control animals (CUMS-sham) into stimulation chambers daily for 1 h per day and connected them to the stimulator without applying any current.

### Drug treatments

Bicuculline (Bic, 2 mg/kg/day; Apexbt, TX, USA) and Sulfobutylether (SBE)-β- Cyclodextrin (10% in normal saline; MCE, China) were administered via intraperitoneal (i.p.) injection for 2 weeks, which was accompanied by the onset of DBS treatment. Mice were kept on drug treatment until the completion of DBS. Control mice received SBE.

### Behavioral studies

All behavioral tests were performed on mice 1 h after the completion of the NAc-DBS or CUMS-sham protocols. The sucrose preference test (SPT), forced swim test (FST), and tail suspension test (TST) were used to assess depression-like behavior [32, 33], whereas general locomotor activity was measured in the open field test (OFT). All behavioral tests were performed according to protocols established in our laboratory. *SPT*. The SPT is widely accepted to measure the extent of anhedonic-like behavior [34, 35]. During the course of the CUMS protocol and DBS treatment, preference for sucrose-sweetened water was measured in mice once a week in their home cages. In mice deprived of food and water for 12 h, exposure to 2% sucrose is used to detect sucrose consumption over 2 h. Bottles were weighed before and after use by experimenters blinded to the mouse group allocation. The sucrose bottle was placed in the position of the original water bottle in the cage. During the ensuing 4 weeks, the intake of 2% sucrose by mice was measured once a week on the day without exposure to stress. The consumption of 2% sucrose solution per unit weight of mice can be calculated according to the following equation: sucrose consumption (g)/body weight (g). Sucrose solution bottles were prepared 24 h before their use. A random sampling of bottles placed in empty cages during SP testing demonstrated that leakage was negligible.

#### FST

The FST was performed in a glass cylinder (2-L beaker) filled with fresh tap water that was maintained at 15°C with an illumination of 100 Lux. Animals were placed individually in the cylinder for 5 min, and their behavior was recorded using a video camera positioned above the basin. The total amount of immobility was scored manually by a trained observer blinded to the treatment groups. An animal was considered immobile when it was floating passively in water, performing only movements that were necessary to keep the nose above the water surface [36].

#### TST

A metal hook was securely fixed with medical adhesive tape to the tip of the tails of the mice, and the animals were suspended using the hook. During the 6 -min testing period, the animals were videotaped and total immobility (reflecting behavioral despair; in which the mice hung without any limb movement) for 5 min (the first and last 30 s were not considered) was scored automatically using analytical software (BIOSEB, In Vivo Research Instruments, France).

#### OFT

The OFT was used to assess anxiety-like behavior [37, 38]. Mice were individually placed into the periphery of the open field (blue plastic box, 50 × 50 × 50 cm; center defined as the inner 25 × 25 cm area; floor illumination, 150 Lux). The overall distance traveled by each mouse was measured using an automated activity monitoring system (DigBehv Video Analysis System, Shanghai Jiliang Software Technology Co., Ltd.) for 5 min.

### Brain microdialysis

The C57BL/6J mice used in this experiment were aged 6–8 weeks. Concentric dialysis probes with a 1.0 mm membrane length were used (CMA 7, Sweden) to detect the concentration of GABA in the VTA. Mice were anesthetized intraperitoneally with pentobarbital sodium (pentobarbital sodium, 0.7%, 10 μL/μg) and placed in a stereotaxic apparatus. The skull was exposed, and a small hole was drilled on one side to expose the dura. The microdialysis catheter was aimed at the unilateral VTA (coordinates: −3.2 mm anterior, 0.4 mm lateral, and 3.0–3.5 mm ventral to bregma). After 5–7 days of recovery, experiments were performed on mice under anesthesia after NAc-DBS. Sterile and oxygenated (95% O_2_, 5 min) artificial cerebrospinal fluid (NaCl, 145 mM; KCl, 3.8 mM; MgCl_2_, 1.2 mM; CaCl_2_, 1.2 mM; pH 7.4) was pumped through the dialysis probe at a constant rate of 1 μL/min (BASi, West Lafayette, USA). A 2-h stabilization period was allowed before the collection of dialysates for analysis, and samples were taken every 30 min from the VTA. Dialysates were automatically collected with a refrigerated autosampler. The three fractions were collected after stimulation. Dialysate samples (15 μL) were injected without purification into the high-performance liquid chromatography-electrochemical detector (HPLC-ECD) apparatus equipped with a chromatographic column (Quattro 3 C18, 3 μm particle size, 150 × 2.1 mm; Chrom4, USA) using a refrigerated autoinjector to quantitate GABA (0.85 V, 35°C). The mobile phase was composed of methanol and buffer salt solution (0.1 M NaH_2_PO_4_; 1:9 (v/v)). This method can only be used after removing bacteria using a 0.22 μm filter and removing bubbles by ultrasonic exposure. Samples or standards were derivatized with phthaldialdehyde (OPA)-Na_2_SO_3_ (37 mM OPA, 50 mM Na_2_SO_3_, 90 mM boric acid, 5% methanol, pH 10.4; 10:1 (v/v)). The mobile phase was pumped at a flow rate of 0.3 mL/min. The retention time for GABA was 30 min. GABA was identified through its migration time and spike profiles. Peak values were normalized to an internal standard curve for a current signal (y, nA × min) and quantified by comparison with an external standard curve (Antec-ALEXYS On-Line Analysis System, Antec®, Netherlands). The values were corrected for in vitro probe recovery, which was approximately 10%.

### Recombinant lentivirus-mediated RNA interference

The lentivirus-mediated interfering RNA (RNAi) was constructed and synthesized by Shanghai GeneChem Co., Ltd. (Shanghai, China). The target sequence used against the mouse GABA A receptor subunit alpha 1 (*GABA_A_-α1*) gene (Gene ID: 14394) was as follows: 5′-TGC CTA ATA AGC TCC TGC GTA-3′. The recombinant lentivirus LV-Gabra1-RNAi_87-GFP (i_87, virus titer: 3 × 10^9^ infectious units mL^−1^) was produced by cotransfecting 293T cells with the lentivirus expression plasmid and packaging plasmids using Lipofectamine 2000. C57BL/6J mice (6–8 weeks of age) were used for stereotactic viral injections. Moreover, lentivirus-mediated RNAi (i_87) knockdown of the *GABA_A_-α1* gene was performed via injection into the bilateral VTA (coordinates: −3.2 mm anterior, ±0.4 mm lateral, and 4.1–4.6 mm ventral to bregma). During pentobarbital sodium anesthesia (as described above), the skull was exposed via a small incision, and a small hole was drilled for injection. A modified microliter syringe (Hamilton) with a 22-gage needle was used: the tip of the needle was placed at the target region, and injection was performed at a speed of 50 nL/min using a micromanipulator. The needle was left in place for 10 min after the injection.

### Neuron-specific adeno-associated viruses stereotaxic injections and optic fiber implantation

The C57BL/6J mice used in this experiment were aged 6–8 weeks. The animals were anesthetized with 0.7% pentobarbital sodium and placed in a stereotaxic frame (RWD, Shenzhen, China). To reveal the type of medium spiny neurons (MSN) projecting to the cell-specific neurons in the VTA, the anterograde monosynaptic transneuronal rAAV-hSyn-Cre-WRPE-PA plasmid (1.0 × 10^13^ infectious units mL^−1^) was injected into the NAc, and a Cre-inducible virus (a double inverted open reading frame (Dio) adeno-associated virus (AAV)) containing the enhanced yellow fluorescent protein (EYFP) (rAAV-Ef1α-Dio-EYFP-WPRE-PA; 1.82 × 10^13^ infectious units mL^−1^) in which the expression of EYFP is driven by the Cre–Dio reaction was injected into the VTA. For calcium imaging in the VTA, a neuron-specific AAV (rAAV2/9-mTH-Gcamp6m-WPRE-pA (5.81 × 10^12^ infectious units mL^−1^; BrainVTA, Wuhan, China)) and rAAV2/9-mDlx-GCaMP6m-WPRE-pA (1.91 × 10^13^ infectious units mL^−1^; BrainVTA, Wuhan, China) were injected into the VTA at two sites (0.4 μL per hemisphere; 0.2 μL at stereotaxic coordinates from bregma: anterior/posterior, −3.2 mm; medial/lateral, ±0.4 mm; dorsal/ventral, −4.1 mm; and 0.2 μL at dorsal/ventral, −4.6 mm, respectively), and an optic fiber was implanted unilaterally at 0.5 mm above the VTA (coordinates: −3.2 mm anterior, ±0.4 mm lateral, and −4.1 mm ventral to bregma). For optogenetic manipulations of the NAc input, we used a combination of a retrograding AAV serotype 2/R carrying Cre-inducible transgenes fused with enhanced green fluorescent protein (eGFP) under the control of the *mDLx* promoter in inhibitory neurons (rAAV2/R-mDLx-Cre-EGFP-WPRE-pA, 4.72 × 10^12^ infectious units mL^−1^) (BrainVTA, Wuhan, China), which was bilaterally injected at two sites of the VTA, to retrogradely trace input neurons to the NAc (0.3 μL per hemisphere; 0.15 μL at stereotaxic coordinates from bregma: anterior/posterior, −3.3 mm; medial/lateral, ±0.5 mm; dorsal/ventral, −4.1 mm; and 0.15 μL at dorsal/ventral, −4.6 mm). Furthermore, a Cre-inducible virus (neuron-specific AAV serotype 2/9 carrying the excitatory optogenetic protein oChIEF and a FLEx element fused to tdTomato under the control of the *CAG* promoter (rAAV2/9-CAG-FLEx-oChIEF-tdTomato-WPRE-pA, 1.54 × 10^13^ infectious units mL^−1^, Taitool Bioscience, Shanghai, China) were bilaterally injected into the NAc (0.2 μL per hemisphere; stereotaxic coordinates from bregma: anterior/posterior, 1.18 mm; medial/lateral, ±1.35 mm; dorsal/ventral, −4.5 mm). For optogenetic manipulations of the VTA input, a neuron-specific AAV serotype 2/9 carrying Cre fused with eGFP under the control of the *NLS* promoter in tyrosine hydroxylase (TH)-positive neurons (rAAV2/9-mTH-NLS-Cre-EGFP-WPRE, 2.34 × 10^12^ infectious units mL^−1^, Taitool Bioscience, Shanghai, China) and a Cre-inducible virus (AAV serotype 2/9 carrying oChIEF and a FLEx element fused to tdTomato under the control of the *CAG* promoter (rAAV2/9-CAG-FLEx-oChIEF-tdTomato-WPRE-pA, 1.54 × 10^13^ infectious units mL^−1^; Taitool Bioscience, Shanghai, China)) were mixed and bilaterally injected at two sites of the VTA (0.3 μL per hemisphere; 0.15 μL at stereotaxic coordinates from bregma: anterior/posterior, −3.3 mm; medial/lateral, ±0.5 mm; dorsal/ventral, −4.1 mm; and 0.15 μL at dorsal/ventral, −4.6 mm). For optogenetic manipulation experiments, mice were also implanted with a bilateral fiber-optic cannula that was secured to the skull using dental cement. Fiber-optic cannulas were 200 μm (fiber core diameter) for optogenetic and fiber photometry experiments. We used the following stereotactic coordinates in the VTA: anterior/posterior, −3.3 mm; medial/lateral, ±1.8 mm; dorsal/ventral, −4.6 mm; 20° angle.

### Optogenetic manipulation

To record calcium imaging in cell-type-specific neurons in the VTA, rAAV2/9-mTH-GCaMP6m and rAAV2/9-mDlx-GCaMP6s were used for injection into the VTA. Fiber photometry allows the real-time excitation and recording of fluorescence from genetic-encoded calcium indicators in freely moving mice. Mice were habituated to the fiber patch cord for at least 15 min per day for 3 days before tests were conducted inside home cages. The fiber photometry system consisted of an excitation light-emitting diode (LED) light (470 nm, Inper Technology Co., Ltd., Hangzhou, China), which was reflected off a dichroic mirror with a 435–488 nm reflection band and a 503–538 nm transmission band (Inper Technology Co., Ltd., Hangzhou, China), and coupled to a 200-μm 0.37 N.A. optical fiber (Inper Technology Co., Ltd., Hangzhou, China) by an objective lens. Laser power was adjusted at the tip of the optical fiber to a low level of 20–30 µW to minimize bleaching. The GCaMP6 fluorescence was bandpass-filtered (MF525-39, Thorlabs) and focused by a 20× objective lens. The fluorescence signal was collected using a CMOS camera. The end of the fiber was imaged at a frame rate of 1–320 Hz. The ROI area size and mean value were set through Inper Studio. Moreover, 470 and 410 nm light sources were used alternately, with 410 nm being used as the control. Analysis of the resulting signal was performed using custom-written MATLAB software. To selectively activate the VTA-GABA neurons projecting MSNs in the NAc, projection-specific expression of oChIEF in the NAc was realized via the combined use of retrograding rAAV2/R-mDLx-Cre-EGFP expression in VTA GABAergic neurons, which was injected into the VTA, and Cre-inducible rAAV-FLEx-oChIEF-tdTomato, which was injected into the NAc. For *in vivo* experiments, the mice were moved to their home cage and allowed a 30 min acclimatization period, the optical cannula was then connected to the optical patch cable, and the mice were administered high-frequency, phasic (frequency: 30 Hz; pulse width: 5 ms; burst duration: 0.267 s; burst interval: 5 s; eight pulses/5 s) blue-light illumination from a blue laser. To selectively activate the DA neurons in the VTA, cell-type-specific expression of oChIEF in dopaminergic neurons was realized via the combined use of rAAV-mTh-Cre-EGFP and Cre-inducible rAAV-FLEx-oChIEF-tdTomato. For *in vivo* experiments, the mice were moved to their home cage and allowed a 30 min acclimatization period, the optical cannula was then connected to the optical patch cable, and the mice were administered high-frequency, phasic (frequency: 20 Hz; pulse width: 40 ms; burst duration: 0.25 s; burst interval: 10 s; five pulses/10 s) blue-light stimulations from a blue laser. In mice expressing oChIEF, a 473 nm blue laser diode and a stimulator were used to generate blue-light pulses. Fibers measuring at least 10 mW were utilized. Phasic light pulses were delivered for 30 min per day for 14 consecutive days. Behavioral testing was resumed at 4 or 24 h poststimulation.

### Histological examination and immunohistochemistry

After completion of all experiments, the mice were anesthetized with pentobarbital sodium intraperitoneally (0.7%, 10 μL/μg) and perfused transcardially with normal saline, followed by 4% paraformaldehyde (PFA). Brains were removed, and coronal sections (40 mm) through the target area were prepared. We confirmed the location of the stimulating electrodes, probes, and optic fibers on bright-field photomicrographs. Only mice with electrodes, probes, and optic fiber placed in the areas of interest were included in subsequent data analyses. Virus-injected brains were fixed overnight in a 4% PFA solution and then equilibrated in 30% sucrose in PBS. Next, 40-μm coronal slices were cut using a freezing microtome. Slices were stored in a cryoprotection solution at 4°C until further processed. The sections were incubated with primary antibodies (mouse anti-GABA_A_ receptor alpha 1 antibody #ab3299, Abcam, 1:1000; rabbit anti-dopamine receptor D1 antibody #ab40653, Abcam, 1:1000; rabbit anti-GABA antibody #ab62669, Abcam, 1:1000; rabbit anti-TH antibody #AB152, Millipore, 1:2000; or mouse anti-Cre recombinase antibody #MAB3120, MERCK, 1:1000) overnight at 4°C. Alexa Fluor®488-, 594-, or 647-conjugated goat antirabbit or antimouse IgG antibodies, or antirabbit IgG antibodies (1:500; Invitrogen, CA, USA) were used as secondary antibodies. Nuclei were counterstained using DAPI. The fluorescent signals were imaged with A1R laser-scanning confocal microscopy using a 10× or 20× objective (Nikon, Japan).

### Western blotting

Brains were immediately flash-frozen after dissection, and defined tissue punches of VTA were then collected using a 1-mm round tissue punch while sectioning brains on a cryostat. Total proteins extracted from the VTA using sodium dodecyl sulfate (SDS) lysis buffer (2% SDS, 10% glycerol, 0.1 mM dithiothreitol, and 0.2 M Tris-HCl, pH 6.8) were subjected to western blotting using the indicated primary (mouse anti-GABA_A_ receptor alpha 1 #ab3299, Abcam, 1:200; rabbit antibeta actin #ab8227, Abcam, 1:1000) and secondary antibodies. Bound antibodies were visualized using enhanced chemiluminescence detection. The band optical densities were quantified by the Image Quant software.

### Data analysis

Data are presented as the mean ± standard error of the mean (SEM). Exact n numbers are given in the table and figure legends. Statistical analysis was performed using GraphPad Prism 7.0 (GraphPad Software, San Diego, CA, USA). Data were statistically analyzed using one-way ANOVA (with post hoc Bonferroni test) or unpaired/paired Student’s *t*-tests. Statistical significance was set at *P* < 0.05. *In vivo* calcium signal analysis/photometry data were exported as MATLAB mat files for further analysis after recording. To visualize the recording traces of the activity of different VTA neuronal populations during DBS_ON, the first 60 s was chosen as the baseline, followed by the calculation of the fluorescence change (*ΔF/F*) normalization by the *ΔF/F* = (*F* − *F_0_*)/*F_0_* method, where *F* is the normalized fiber photometry signal data and *F_0_* is the average of the fluorescence values during the baseline period (from 60 s preceding the time of DBS_ON). The *ΔF/F* values are presented with heatmaps or per-event plots, with shaded areas indicating the SEM. All calculations were performed using MATLAB 2014a.

## Results

### NAc**-DBS reduced depression-like behavior in CUMS mice**

Two weeks after the CUMS model was established, mice implanted with stimulation electrodes were monitored for depressive-like manifestations using a battery of behavioral tests (Figures 1A and B). Compared with wild-type (WT) mice, CUMS-induced a reduction in sucrose consumption (Figure 1C) and an increase in the time spent immobile both in the TST (Figure 1D) and FST (Figure 1E). In addition, a decrease in open field behavior was observed in CUMS mice, including a reduction in the total distance, central distance, and time spent in the center, as well as an increase in the resting time in the OFT (Figure 1F). These results suggest that after the 2-week CUMS regimen, the susceptible mice gradually developed a decrease in reward sensitivity or anhedonia and an increase in the level of despair and helplessness [31]. We next examined whether the long-term effects of high-frequency-dependent NAc-DBS would alter the depression-like behavior of CUMS mice. A single session of high-frequency-dependent NAc-DBS or long-term, low-frequency (<50 Hz) stimulation had no effect and resulted in unaltered depressive symptoms in CUMS mice, such as anhedonia (data not shown). In fact, we found that DBS did not effectively improve the depressive-like behavior of CUMS mice after 1 week of continuous stimulation, including both sucrose consumption and TST (Figures 1C and D). By contrast, CUMS mice that received repeated (2 weeks) NAc-DBS (100 Hz) displayed a significant increase in sucrose consumption and locomotor activity, which returned to the normal level compared with WT mice. Moreover, the reduction in the amount of time spent immobile both in the TST and FST was reversed by the DBS treatment (Figures 1C–F). Our results indicate that chronic and high-frequency-dependent NAc-DBS has a robust antidepressant effect.

**Figure 1.**
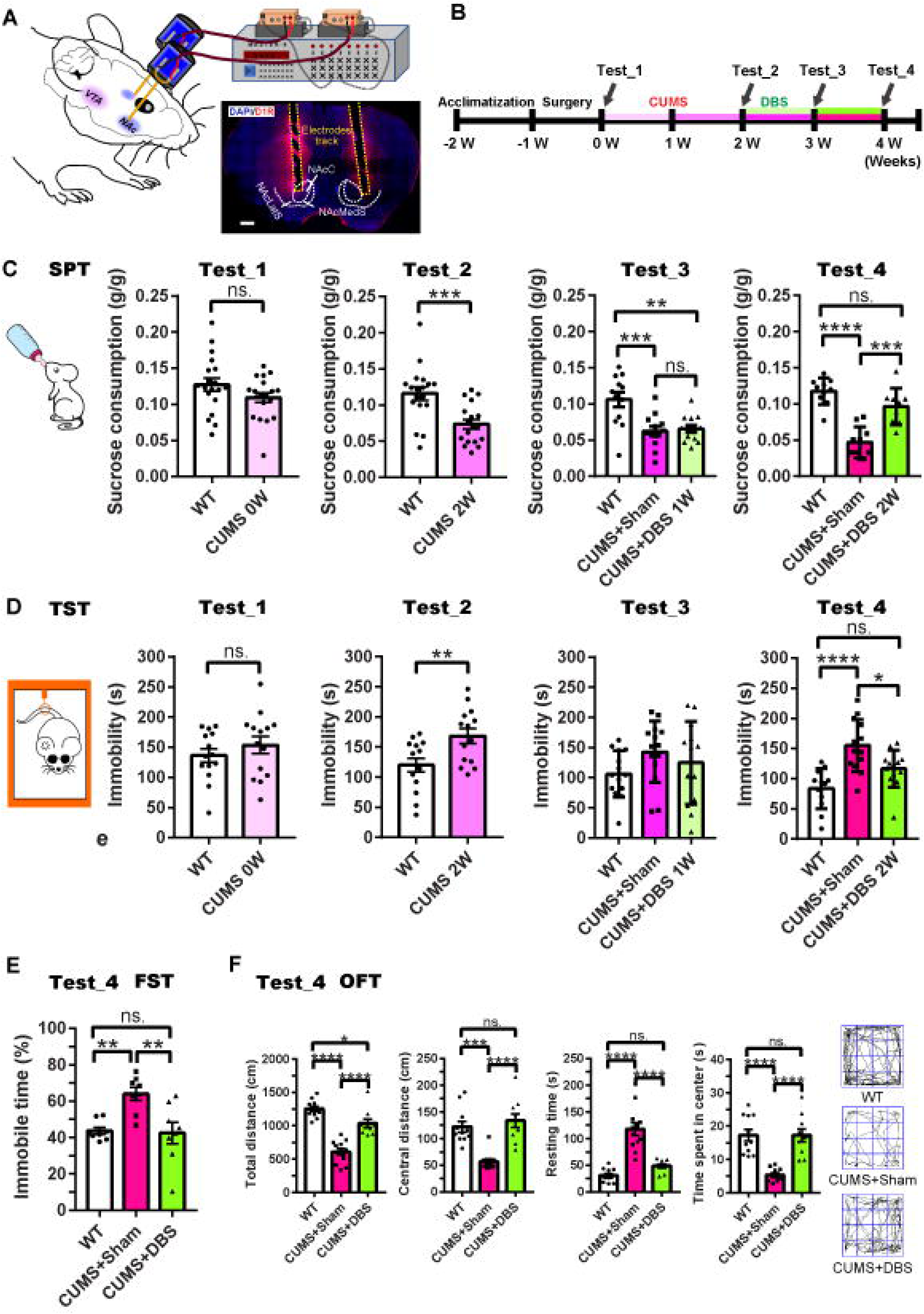
Long-term and high-frequency-dependent nucleus accumbens deep brain stimulation (NAc-DBS) attenuated depression-like behaviors in chronic unpredictable mild stress (CUMS) mice. (A) Experimental setup and schematic diagram of electrode placement for DBS. The white dotted line represents the NAc area, and the yellow dotted line indicates the electrode tracks (NacC, nucleus accumbens core; NacLatS, lateral shell area of the nucleus accumbens; NacMedS, medial shell area of the nucleus accumbens; scale bar, 500 μm). (B) Timeline of electrode implantation and the battery of behavioral tests used here. The red and blue lines represent the CUMS and deep brain stimulation (DBS) periods, respectively. (C) The sucrose preference test (SPT) was used to measure the extent of anhedonic-like behavior during the course of the CUMS protocol and DBS treatment every week (unpaired *t*-test: CUMS 0W, *t* = 1.586, *P* = 0.1216, n = 19 animals per group; CUMS 2W, *t* = 3.765, *P* = 0.0006, n = 18 animals per group; one-way ANOVA: CUMS 3W/DBS 1W, F_(2, 35)_ = 10.65, *P* = 0.0002, n = 12–13 animals per group; CUMS 4W/DBS 2W, F_(2, 26)_ = 28.09, *P* < 0.0001, n = 10 animals per group). Data represent the means ± SEM; \*\**P* < 0.01, \*\*\**P* < 0.001, \*\*\*\**P* < 0.0001. (D) The tail suspension test (TST) was used to assess depression-like behavior during the course of the CUMS protocol and DBS treatment every week (unpaired *t*-test: CUMS 0W, *t* = 0.9479, *P* = 0.3519, n = 14 animals per group; CUMS 2W, *t* = 2.861, *P* = 0.0082, n = 14 animals per group; one-way ANOVA: CUMS 3W/DBS 1W, F_(2, 38)_ = 1.624, *P* = 0.2104, n = 14 animals per group; CUMS 4W/DBS 2W, F_(2, 35)_ = 12.18, *P* < 0.0001, n = 12–13 animals per group). Data represent the means ± SEM; \**P* < 0.05, \*\**P* < 0.01, \*\*\*\**P* < 0.0001. (E) The FST was used to assess depression-like behavior after 2 weeks of NAc-DBS. Relative to their WT counterparts, mice in the CUMS+Sham group displayed an enhanced depression-like behavior, as indicated by an increased period of immobility in the forced swim test. Chronic treatment with NAc-DBS rescued the depressive phenotype of CUMS mice. Data represent the means ± SEM; \*\**P* < 0.01, \*\*\**P* < 0.001, \*\*\*\**P* < 0.0001, one-way ANOVA (F_(2, 21)_ = 8.582, *P* = 0.0019, n = 8 animals per group). (F) The OFT was used to measure locomotor activity after 2 weeks of NAc-DBS. The left panels are the total distance (F_(2, 30)_ = 44.84, *P* < 0.0001, n = 11–12 animals per group), central distance (F_(2, 28)_ = 17.31, *P* < 0.0001, n = 10–11 animals per group), resting time (F_(2, 31)_ = 36.81, *P* < 0.0001, n = 12 animals per group), and time spent in the center (F_(2, 28)_ = 19.09, *P* < 0.0001, n = 10–11 animals per group), respectively. The right panels are the trajectory maps in the OFT of a typical mouse from each group. Data represent the means ± SEM; \*\**P* < 0.01, \*\*\**P* < 0.001, \*\*\*\**P* < 0.0001, one-way ANOVA.

### A single trial of NAc-DBS increased both the level of the GABA neurotransmitter and dopaminergic neuron activity in the VTA

According to literature [39, 40] and the previous experimental verification performed in our laboratory, we found that most of the Cre-expressed cells in the NAc were dopamine receptor type 1 (D1R) positive neurons (Supplementary Figures 1A and B), and there is a synaptic connection between the NAc and VTA (Supplementary Figure 1C). To assess the effect of NAc-DBS on the GABA neurotransmitter levels in the VTA, a single trial was conducted. The experimental process is shown in Figure 2A. Within 30 min of the first sample collection (i.e., Sample_1 at Time_1), a significant increase in the level of the GABA neurotransmitter in the VTA of the WT mice that underwent stimulation treatment once (i.e., WT+DBS) was observed compared with the control (i.e., WT+DBS) (Figure 2B). However, there was no significant difference in GABA neurotransmitter levels between the two groups during the second 30 min of the next sample collection (i.e., Sample_2 at Time_2) (Figure 2B). We also found that in both the control mice and the NAc-DBS treatment mice, the levels of the GABA neurotransmitter between the two samples (i.e., Sample_1 and Sample_2) in the same group were significantly reduced after 30 min (Figure 2C). These results demonstrated that acute electrical stimulation of NAc can significantly increase the level of the GABA neurotransmitter in the VTA of WT mice but this increased GABA will be decreased dramatically within a short time.

**Figure 2.**
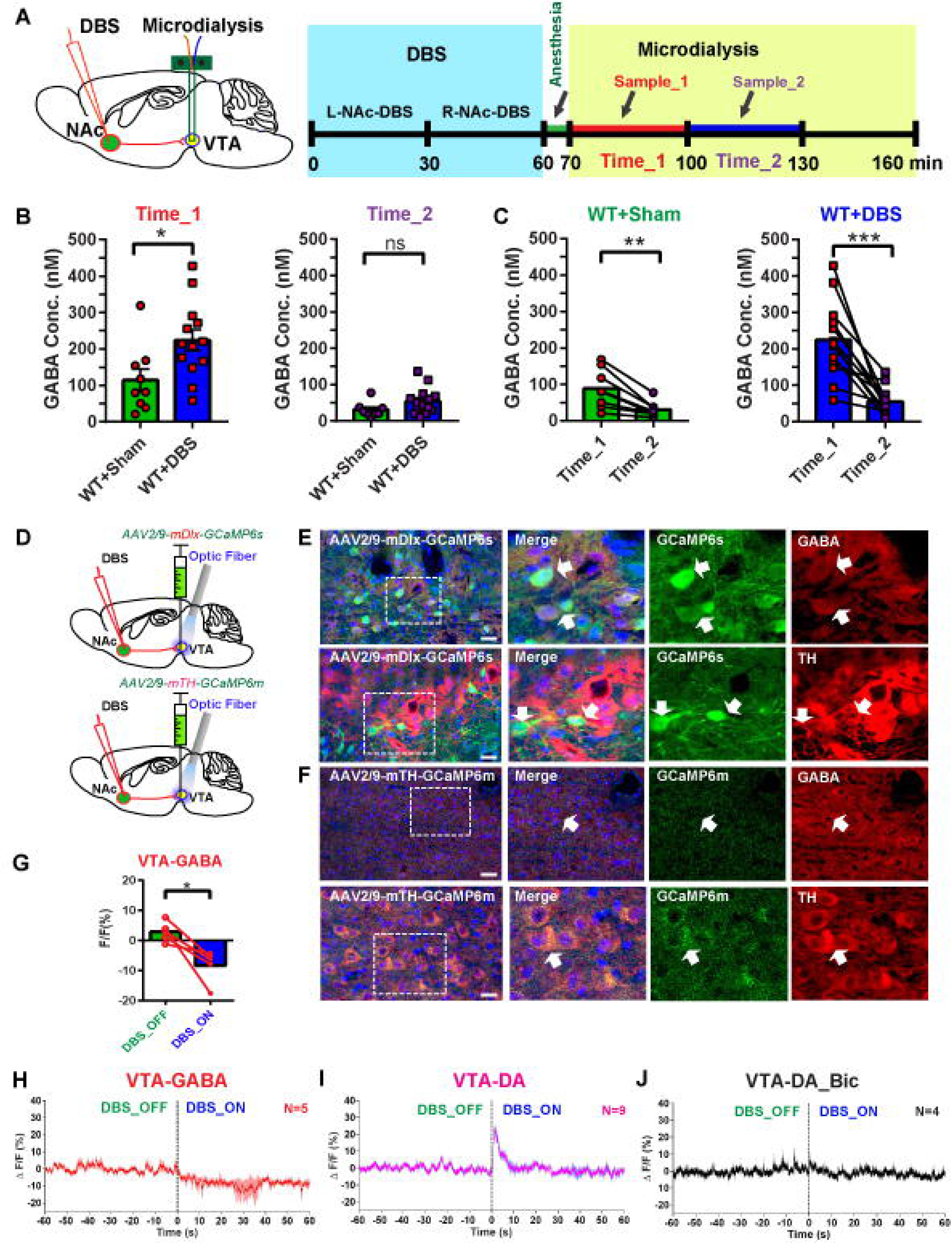
A single trial of high-frequency-dependent nucleus accumbens deep brain stimulation (NAc-DBS) increased both the level of the γ-aminobutyric acid (GABA) neurotransmitter and dopaminergic neuron activity in the ventral tegmental area (VTA). (A) The panel on the left is the experimental setup: stimulation electrodes were implanted into the bilateral NAc, and a microdialysis catheter with a probe was embedded into the unilateral VTA. The panel on the right is the timeline of the acute electrical stimulation of the NAc and the microdialysis sample collection (L-NAc-DBS: the left NAc was stimulated; R-NAc-DBS: the right NAc was stimulated). (B) Concentrations of the GABA neurotransmitter in Sample_1 and Sample_2 between mice in the WT+Sham group and mice in the WT+DBS group during Time_1 and Time_2, respectively (unpaired *t*-test: Time_1, *t* = 2.537, *P* = 0.0196, n = 9 in the WT+Sham group, n = 13 in the WT+DBS group; Time_2, *t* = 1.677, *P* = 0.1091, n = 8 in the WT+Sham group, n = 14 in the WT+DBS group). Data represent the means ± SEM; \**P* < 0.05. (C) Concentrations of the GABA neurotransmitter in Sample_1 and Sample_2 in the WT+Sham group during Time_1 and Time_2, respectively, and concentrations of the GABA neurotransmitter of Sample_1 and Sample_2 in the WT+DBS group during Time_1 and Time_2, respectively (paired *t*-test: WT+Sham, *t* = 4.422, *P* = 0.0031, n = 8 animals per group; WT+DBS, *t* = 5.549, *P* = 0.0001, n = 13 animals per group). (D) Experimental setup: stimulation electrodes were implanted into the bilateral NAc, and cell-specific rAAV-GCaMP6 was injected into the bilateral VTA, which was stimulated *in vivo* by illuminating the unilateral VTA. (E) and (F) Confocal images of VTA slices from VTA-GABA (GABA^+^) neurons (red) and VTA-DA (TH^+^) neurons (red) injected intra-VTA with rAAV2/9-mDlx-GCaMP6s (E) or rAAV2/9- mTH-GCaMP6m (green), respectively. Scale bar, 500 μm. (G)–(I) Representative traces demonstrating the temporal profile of VTA Ca^2+^ activity from different neuronal populations during DBS_OFF and DBS_ON, including VTA-GABA neurons (G: paired *t*-test, *t* = 3.68, *P* = 0.0212, n = 5 animals per group; and H) and VTA-DA neurons (I), as well as VTA-DA neurons of mice injected with Bic in advance (J). Data are presented as the percent change in fluorescence over the mean fluorescence (ΔF/F), n = 4–9 animals per group.

Next, we introduced the cell-specific expressed CaMP6 protein in the VTA of WT mice (Figure 2D). Subsequently, we confirmed the cell-specific expression of CaMP6 in VTA-GABA and VTA-DA neurons via an immunohistochemical method. Our results showed that green fluorescent protein (GFP)-positive cells were never observed outside of the VTA. Only a small number of GFP-positive and GABA-negative or GFP-positive and TH-negative cells were observed among all GFP-positive cells in the VTA. Most of the GABA-positive cells or TH-positive cells were also GFP-positive (Figures 2E and F). Calcium imaging experiments were used to record the activity of specific neuronal populations in the VTA, such as GABAergic neurons (i.e., VTA-GABA) and dopaminergic neurons (i.e., VTA-DA). Using fiber photometry, we appraised the real-time activity of VTA-GABA and VTA-DA neurons in awake mice in response to acute NAc-DBS treatment (Figures 2G–I) or bicuculline (Bic, a GABA_A_ receptor antagonist [24], Figure 2J). In VTA-GABA neurons, GCaMP6 fluorescence intensity exhibited a sustained and slow decrease, i.e., there was an apparent change in the activity of VTA-GABA neurons during the DBS_ON period (0%–10% Δ*F*/*F*, Figures 2G and H). However, soon after the onset of NAc-DBS, the CaMP6 fluorescence rapidly increased over the range of basal fluctuation and reached its peak (30%–40% Δ*F*/*F*) at 0–1 s in the VTA-DA neurons. The increased fluorescence of CaMP6 expressed in VTA-DA neurons rapidly disappeared within 3–6 s, indicating a rapid and transient neuronal activation (Figure 2I). Interestingly, we also found that the rapid increase in GCaMP6 fluorescence intensity in VTA-DA neurons, which induced NAc-DBS onset, could be completely blocked by Bic (Figure 2J).

### Chronic optical stimulation of VTA-projecting NAc neurons alleviates the depression-like behavior of CUMS mice

To determine whether the direct NAc–VTA pathway was critical for the antidepressant effect of NAc-DBS, we specifically labeled VTA_DA (Th^+^) neurons via bilateral injection of Cre-dependent AAV2/9-CAG-FLEx-oChIEF-tdTomato and AAV2/9-mTH-NLS-Cre-EGFP into the VTA for the expression of oChIEF in VTA_DA (Th^+^) neurons and implanted optical fibers in the VTA. Repeated optogenetic stimulation paradigms altered emotional behaviors upon 2 weeks of illumination of oChIEF-expressing VTA dopamine neurons in the VTA (LED_VTA-DA, Supplementary Figures 2A–C), including ameliorated sucrose consumption in the SPT (Supplementary Figure 2D), immobility time in the TST (Supplementary Figure 2E), and locomotor activity in the OFT (Supplementary Figure 2F). Moreover, we specifically expressed oChIEF in the VTA_GABA-projecting NAc neurons through bilateral injections of AAV2/retro-mDLx-Cre expressing retrogradely transported Cre recombinase into the GABA neurons of the VTA and AAV2/9-CAG-FLEx-oChIEF-tdTomato expressing Cre-dependent oChIEF into the NAc. Subsequently, we implanted optical fibers in the VTA (Figures 3A and B). Repeated specific activation of VTA-GABA-projecting MSN terminals of NAc (LED_VTA-GABA) also exerted antidepressant effects in CUMS mice (Figure 3). Optogenetics of the NAc–VTA pathway is sufficient to induce an antidepressant effect similar to that of DBS.

**Figure 3.**
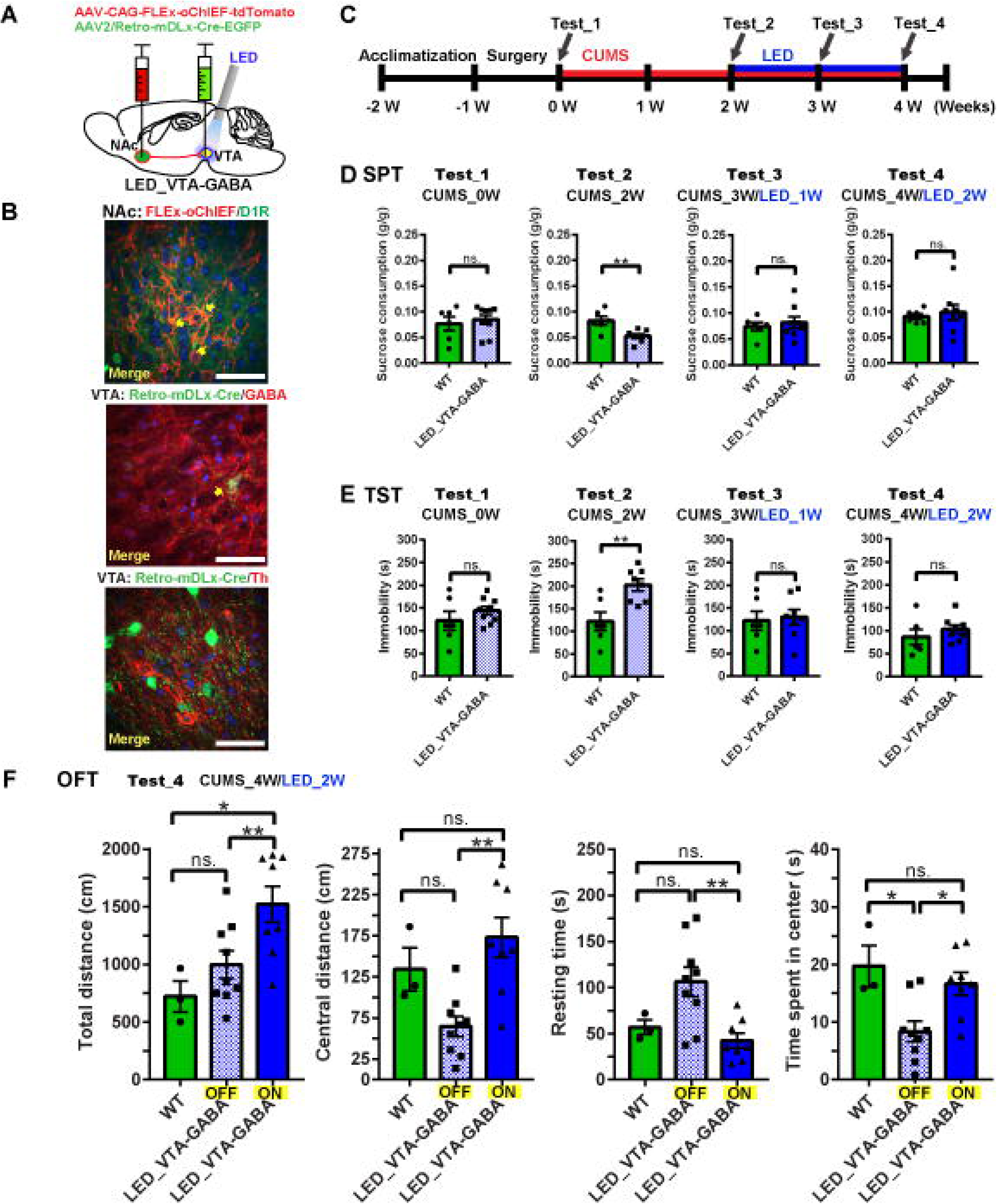
Chronic optical stimulation of VTA-GABA-projecting MSN terminals of NAc alleviates the depression-like behavior in CUMS mice. (A) Schematic representation of the oChIEF-mediated activation of VTA-GABA neurons in CUMS mice. (B) Upper panel: representative image showing oChIEF-tdTomato-labeled VTA_GABA- projecting NAc neurons; middle panel: representative image showing mDLx-Cre- EGFP-labeled VTA_GABA neurons (GABA^+^); bottom panel: representative image showing that Cre-EGFP did not colabel with VTA_DA neurons (Th^+^). Scale bar, 50 μm. (C) Timeline of the virus injection and battery of behavioral tests performed here. The red and blue lines represent the CUMS and chronic optical stimulation periods, respectively. (D) The sucrose preference test (SPT) was used to measure the extent of anhedonic-like behavior during the course of the CUMS protocol and optical stimulation treatment every week (CUMS_0W, WT, 0.07664 ± 0.01326 g/g, n = 6; LED_VTA-GABA, 0.08493 ± 0.009056 g/g, n = 9; *t* = 0.5363, *P* = 0.6008; CUMS_2W, WT, 0.08262 ± 0.008272 g/g, n = 6; LED_VTA-GABA, 0.05242 ± 0.004139 g/g, n = 8; *t* = 3.53, *P* = 0.0041; CUMS_3W/LED_1W, WT, 0.07386 ± 0.00792 g/g, n = 6; LED_VTA-GABA, 0.08181 ± 0.01127 g/g, n = 8; *t* = 0.5376, *P* = 0.6007; CUMS_4W/LED_2W, WT, 0.09075 ± 0.005518 g/g, n = 6; LED_VTA-GABA, 0.09882 ± 0.01494 g/g, n = 8; *t* = 0.447, *P* = 0.6628). (E) The tail suspension test (TST) was used to assess the depression-like behavior during the course of the CUMS protocol and optical stimulation treatment every week (CUMS_0W, WT, 121.9 ± 20.82 s, n = 6; LED_VTA-GABA, 144.2 ± 9.37 s, n = 9; *t* = 1.097, *P* = 0.2925; CUMS_2W, WT, 121.9 ± 20.82 s, n = 6; LED_VTA-GABA, 202.4 ± 12.92 s, n = 8; *t* = 3.454, *P* = 0.0048; CUMS_3W/LED_1W, WT, 121.9 ± 20.82 s, n = 6; LED_VTA-GABA, 129.5 ± 16.73 s, n = 8; *t* = 0.2881, *P* = 0.7782; CUMS_4W/LED_2W, WT, 86.3 ± 16.31 s, n = 6; LED_VTA-GABA, 103 ± 9.579 s, n = 8; *t* = 0.9332, *P* = 0.3691). (F) The open field test was used to measure locomotor activity after 2 weeks of optical stimulation (total distance, F_(2, 17)_ = 6.315, *P* = 0.0089, n = 3–9 animals per group; central distance, F_(2, 17)_ = 9.147, *P* = 0.0020, n = 3–9 animals per group; resting time, F_(2, 17)_ = 6.923, *P* = 0.0063, n = 3–9 animals per group; and time spent in the center, F_(2, 17)_ = 6.842, *P* = 0.0066, n = 3–9 animals per group). The data in D and E represent the means ± SEM; \*\**P* < 0.01, \*\*\**P* < 0.001, \*\*\*\**P* < 0.0001, *t*-test. The data represent the means ± SEM; \*\**P* < 0.01, \*\*\**P* < 0.001, \*\*\*\**P* < 0.0001, one-way ANOVA.

### GABA_A_ receptor inhibition attenuates the effect of NAc-DBS

In the present work, we evaluated the NAc-to-VTA connectivity to determine whether VTA dopaminergic neurons are modulated by NAc stimulation and determine the role of GABA and GABA_A_ receptors in the response of VTA to NAc stimulation. Thus, CUMS mice were administrated bicuculline (Bic, 2 mg/kg/day, intraperitoneal injection) continuously. According to the timeline of electrode implantation and pharmacological intervention provided in Figure 4A, compared with the solvent control group (i.e., CUMS+SBE+DBS), CUMS mice treated with DBS after Bic administration (i.e., CUMS+Bic+DBS) did not exhibit an effective improvement in their depressive-like behavior after 2 weeks of continuous stimulation, including both a reduction in sucrose consumption (Figure 4B) and an increase in the total amount of immobility time in the TST (Figure 4C). In addition, Bic did not improve the locomotor activity of CUMS mice treated with NAc-DBS in the OFT (Figure 4D). In turn, mice in the CUMS+Bic+DBS group had depression-like behavior phenotypes that were similar to those of the CUMS mouse model (i.e., CUMS+Bic+Sham). However, Bic did not affect a series of behavioral modifications in the normal WT (i.e., WT+Bic) compared with the WT group. The results showed that systemic administration of Bic blocked the antidepressant effect of NAc-DBS by inhibiting GABA_A_ receptors.

**Figure 4.**
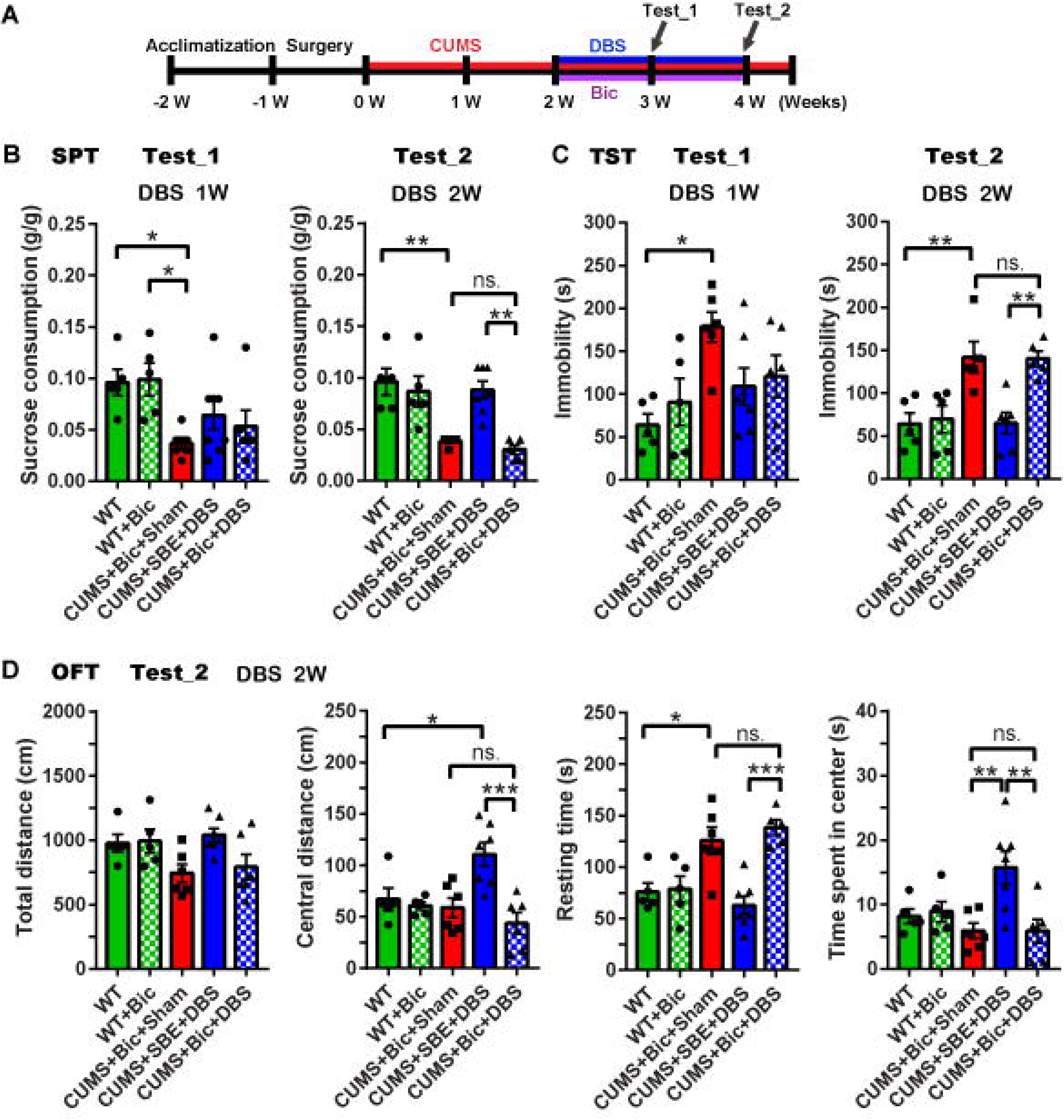
Pharmacological characterization of bicuculline showing that it can block the antidepressant effect of long-term and high-frequency-dependent nucleus accumbens deep brain stimulation (NAc-DBS). (A) Timeline of the experimental procedure used in this study. The red and blue lines represent the CUMS and deep brain stimulation (DBS) periods, respectively, whereas the purple line represents bicuculline administration (2 mg/kg per day, i.p.). (B) The sucrose preference test (SPT) was used to measure anhedonic-like behavior during the course of the CUMS protocol accompanied by DBS treatment over 2 weeks. The data represent the means ± SEM; \**P* < 0.05, one-way ANOVA (CUMS 3W/DBS 1W: F_(4, 25)_ = 3.62, *P* = 0.0185, n = 5–8 animals per group; CUMS 4W/DBS 2W: F_(4, 23)_ = 10.63, *P* < 0.0001, n = 5–7 animals per group). (C) The tail suspension test (TST) was used to assess depression-like behavior during the course of the CUMS protocol accompanied by DBS treatment over 2 weeks. The data represent the means ± SEM; \*\**P* < 0.01, one-way ANOVA (CUMS 3W/DBS 1W: F_(4, 24)_ = 3.713, *P* = 0.0173, n = 5–7 animals per group; CUMS 4W/DBS 2W: F_(4, 21)_ = 8.428, *P* = 0.0003, n = 5–6 animals per group). (D) The open field test was used to measure locomotor activity after 2 weeks of NAc-DBS accompanied by bicuculline administration. The data represent the means ± SEM; \**P* < 0.05, \*\**P* < 0.01, \*\*\**P* < 0.001, one-way ANOVA (total distance: F_(4, 24)_ = 3.159, *P* = 0.0321, n = 5–7 animals per group; central distance: F_(4, 24)_ = 7.191, *P* = 0.0006, n = 5–7 animals per group; resting time: F_(4, 23)_ = 10.24, *P* < 0.0001, n = 5–7 animals per group; time spent in the center: F_(4, 24)_ = 5.397, *P* = 0.0030, n = 5–7 animals per group).

To determine whether NAc-DBS increased the activity of VTA-DA neurons by decreasing the activity of VTA GABAergic interneurons, we used lentivirus-mediated RNAi to knock down the expression of the GABA_A_ receptor subunit alpha 1 (*GABA_A_-α1*) gene in VTA-GABA neurons but not in dopaminergic neurons [41, 42]. We found that blocking GABA_A_-α1 reversed the DBS induced behavioral modifications (Figure 5). The positions of the lentivirus (LV-RNAi_87-GFP) injection and electrode implantation, as well as the experimental procedure, are shown in Figures 5A and B, respectively. LV-RNAi_87-GFP led to the knockdown of the target mouse *GABA_A_-α1* gene expression in the VTA (Figure 5C). First, the lentivirus injection did not affect the establishment of the CUMS mouse model in each group, such as the reduction in sucrose consumption (Supplementary Figure 3A) and increase in the total amount of immobility time in the TST (Supplementary Figure 3B). These changes developed gradually over time of CUMS exposure. However, in the 2-week continuous NAc-DBS intervention, CUMS mice injected with LV-RNAi_87-GFP (i.e., CUMS+i_87+DBS) in advance did not exhibit an improvement in their depressive symptoms after high-frequency stimulation, i.e., the sucrose consumption (Figure 5D), immobility time in the TST (Figure 5E), and locomotor activity in the OFT (Figure 5F) could not be reversed by NAc-DBS compared with the CUMS mouse model group (i.e., CUMS+i_87+Sham). Moreover, compared with CUMS mice injected with the control lentivirus (LV-RNAi_vehicle-GFP; i.e., CUMS+i_vehicle+Sham), 2 weeks of DBS effectively ameliorated a series of depressive-like behavioral phenotypes in CUMS mice (i.e., CUMS+i_vehicle+DBS) but obviously did not improve those of mice in the CUMS+i_87+DBS group, which exhibited a persistent depression-like behavior, such as performance in the SPT (Figure 5D), TST (Figure 5E), and OFT (Figure 5F). These results indicated that the cell-specific knockdown of the *GABA_A_-α1* gene and consequent downregulation of GABA_A_-α1 expression in VTA_GABA neurons blocked the antidepressant effect of DBS on the CUMS-induced depressive phenotype.

**Figure 5.**
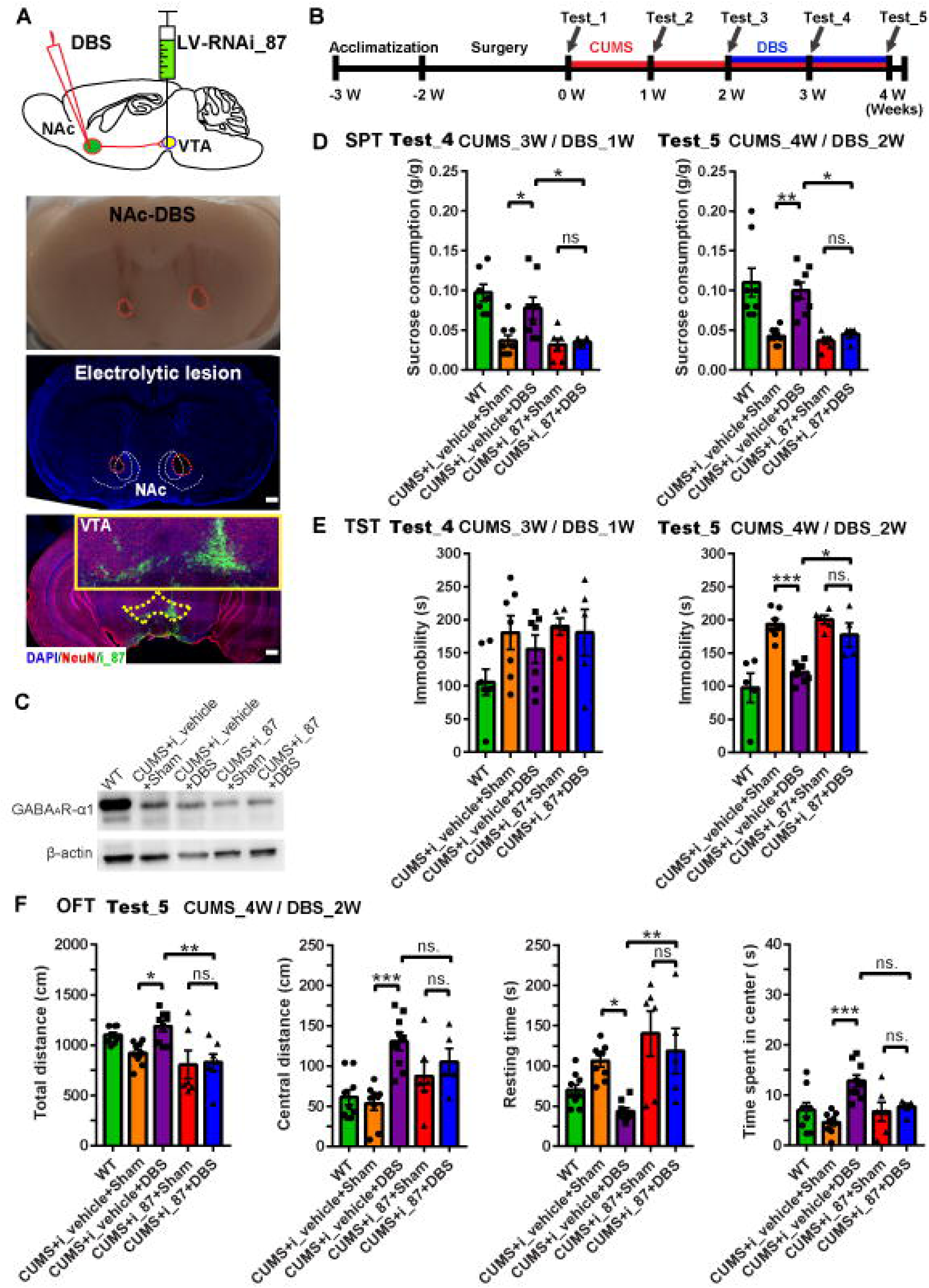
Knockdown of γ-aminobutyric acid A subunit alpha 1 (GABA_A_-α1) receptors blocked the antidepressant effect of long-term and high-frequency-dependent nucleus accumbens deep brain stimulation (NAc-DBS). (A) The top panel illustrates the experimental setup: stimulation electrodes were implanted into the bilateral NAc, and LV_RNAi_87 was injected into the bilateral VTA. The middle two images correspond to a bright-field photograph and a confocal image of coronal sections of the NAc after DBS electrical damage. The red dotted circle represents the lesion area of the NAc created by a strong current (300 μA), and the white dotted circle represents the NAc area. The bottom panel corresponds to confocal images of VTA slices from VTA neurons injected with LV_RNAi_87 (green). Scale bar, 500 μm. (B) Timeline of the experimental procedure used in this study. The red and blue lines represent the CUMS and deep brain stimulation (DBS) periods, respectively. (C) Western blot detection of GABA_A_-α1 receptors in the different treatment groups. (D) The sucrose preference test (SPT) was used to measure anhedonic-like behavior during the course of the CUMS protocol accompanied by 2 weeks of DBS treatment in mice injected with i_87 into the VTA in advance. The data represent the means ± SEM; \**P* < 0.05, \*\**P* < 0.01, one-way ANOVA (CUMS 3W/DBS 1W: F_(4, 31)_ = 9.479, *P* < 0.0001, n = 6–9 animals per group; CUMS 4W/DBS 2W: F_(4, 29)_ = 9.889, *P* < 0.0001, n = 4–8 animals per group). (E) The tail suspension test (TST) was used to assess depression-like behavior during the course of the CUMS protocol accompanied by 2 weeks of DBS treatment in mice injected with i_87 into the VTA in advance. The data represent the means ± SEM; \**P* < 0.05, \*\*\**P* < 0.001, one-way ANOVA (CUMS 3W/DBS 1W: F_(4, 26)_ = 2.178, *P* = 0.0996, n = 5–7 animals per group; CUMS 4W/DBS 2W: F_(4, 24)_ = 15.1, *P* < 0.0001, n = 4–8 animals per group). (F) The open field test was used to measure locomotor activity after 2 weeks of NAc-DBS in mice injected with i_87 into the VTA in advance. The data represent the means ± SEM; \**P* < 0.05, \*\**P* < 0.01, \*\*\**P* < 0.001, one-way ANOVA (total distance: F_(4, 35)_ = 6.368, *P* = 0.0006, n = 6–9 animals per group; central distance: F_(4, 33)_ = 8.121, *P* = 0.0001, n = 5–9 animals per group; resting time: F_(4, 33)_ = 7.748, *P* = 0.0002, n = 5–9 animals per group; time spent in the center: F_(4, 33)_ = 6.896, *P* = 0.0004, n = 5–9 animals per group).

On the basis of these findings, we hypothesized that GABA_A_-α1 receptors play an important role in the connectivity induced by DBS.

### GABA_A_-α1 receptors in VTA-GABA neurons mediate the process of NAc-DBS disinhibition of VTA-DA neurons

To further clarify whether the effect of NAc-DBS on the correction of depression-like behaviors in the CUMS mouse model was mediated by VTA-GABA neuron disinhibition to DA neurons in the VTA, we used lentivirus-mediated RNAi (LV-RNAi_87) to knock down the expression of the GABA_A_ α1 subunit in VTA-GABA neurons, followed by the injection of rAAV2/9-mTh-GCaMP6m (which is expressed specifically in VTA-DA neurons) and the implantation of optical fibers in the VTA (Figure 6A). Consistent with existing literature, expression of GABA_A_-α1 was observed mainly in VTA-GABA neurons, and LV-RNAi_87 knocked down GABA_A_-α1 in the VTA_GABA neurons exclusively (Figure 6B and Supplementary Figure 4). *In vivo* fiber photometry Ca^2+^ imaging analysis revealed that the knockdown of the *GABA_A_-α1* gene in VTA-GABA neurons led to the inability of NAc-DBS to increase the fluorescence signals of GCaMP6 in VTA-DA neurons, as shown in Figure 2H (Figures 6C and D). However, the tail suspension stress increased the fluorescence signals of GCaMP6 in VTA-DA neurons (Figures 6E and F). These results suggest that knockdown of the *GABA_A_-α1* gene in VTA-GABA neurons can block the effect of NAc-DBS on the increase in VTA-DA neuronal activity. Moreover, in the absence of NAc-DBS, *GABA_A_-α1* gene knockdown in VTA-GABA neurons did not affect the functional activity of DA neurons.

**Figure 6.**
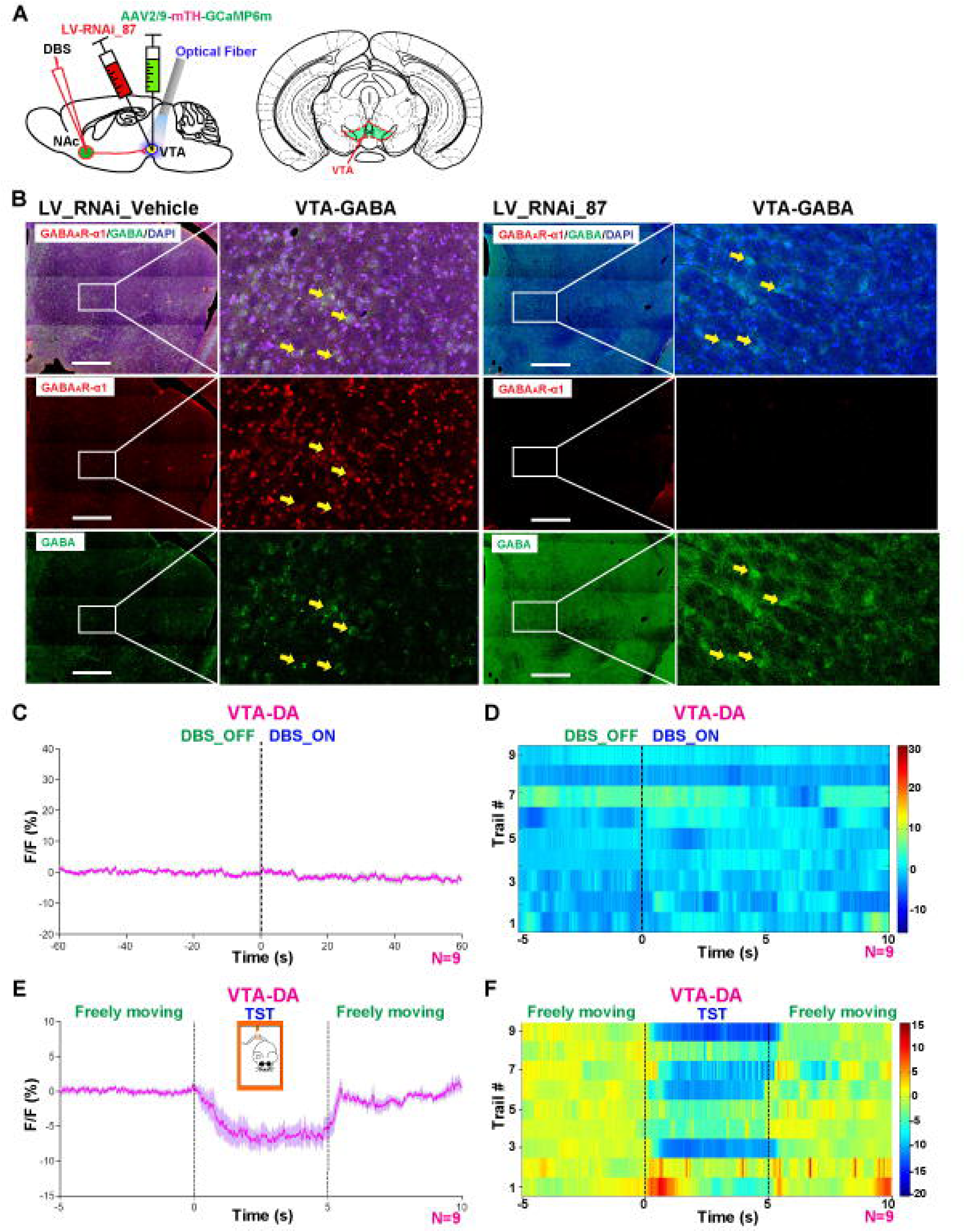
GABA_A_-α1 receptors in VTA-GABA neurons mediate the process of NAc- DBS disinhibition of VTA-DA neurons. (A) Schematic representation of the RNAi- mediated inactivation of VTA-DA neurons in mice with NAc-DBS treatment. (B) Confocal images of VTA slices from VTA neurons injected with LV_RNAi_87 or LV_RNAi_Vehicle. VTA_GABA (GABA^+^) neurons (green) colabeled with an anti- GABA_A_-α1 antibody (red) in mice injected with LV_RNAi_Vehicle. However, VTA_GABA (GABA^+^) neurons (green) were not colabeled with the anti-GABA_A_-α1 antibody (red) in mice injected with LV_RNAi_87. Scale bar, 500 μm. (C) and (E) Representative traces demonstrating the temporal profile of the VTA_DA neuron Ca^2+^ activity during DBS_OFF/DBS_ON (C) and tail suspension stress (E). (D) and (F) Representative heat map of the Ca^2+^ signal activity recorded in each mouse during DBS_OFF/DBS_ON (D) and tail suspension stress (F). Data are presented as the percent change in fluorescence signals over the mean fluorescence (ΔF/F), n = 9.

Taken together, these data suggest that the acute onset of NAc stimuli results in an increase in the levels of the GABA neurotransmitter, accompanied by the increase in VTA-DA neuronal activity in the VTA. Finally, GABA and its GABA_A_ receptors seem to be involved in the above-mentioned neurobiological process of NAc-DBS onset.

## Discussion

DBS involves the implantation of electrodes within specific brain regions that provide electrical stimulation to the brain to create a therapeutic effect [43]. DBS is effective in relieving the symptoms of movement disorders, most notably Parkinson’s disease and essential tremor [44]. In addition, depressive symptoms in patients with TRD can now be alleviated over a period of weeks by NAc-DBS at a high-frequency (145 Hz) [13], and long-term treatment appears to be an important factor for significantly reducing the depressive symptoms of these patients compared with acute stimulation [7, 8, 34, 45]. Similarly, in a mouse model of depression, repeated high-frequency stimulation produced a robust antidepressant effect, whereas acute stimulation was ineffective [6]. However, the investigation of the mechanisms underlying these phenomena is limited in humans, for understandable reasons [6], and preclinical data showing the behavioral efficacy of DBS, as well as its neural circuit excitability modulation correlates, are rare in relevant animal models.

Converging human and rodent studies demonstrated a critical role of the NAc in depression symptomatology, including reduced motivation and anhedonia [46–49]. The NAc integrates information from afferent inputs leading to motivation and reward functions [50, 51]. Recent studies demonstrated that these NAc afferent inputs become dysfunctional after exposure to stressful stimuli, resulting in altered cellular and molecular mechanisms in the NAc that mediate depression-like outcomes [34]. NAc D1-MSNs project to the ventral pallidum, internal globus pallidus, ventral tegmental area (VTA), and substantia nigra, whereas NAc D2-MSNs project to the ventral pallidum [52, 53]. These two neuronal populations work together to promote normal behavioral outputs, whereas imbalance in one MSN subtype can trigger dysfunctional motivational states [54, 55]. This network balance model is supported by positive and negative outcome tasks in humans and by studies in rodents, which demonstrate a role for the activation of the D1-MSN pathway in positive reward and activation of the D2-MSN pathway in aversion, whereas inhibition of these pathways produces opposing outcomes [34, 56–58]. Zhu et al. reported that in contrast with control mice, GABAergic neurons in the NAc from mice with CUMS-induced depression are featured as decreases in inhibitory synapse outputs (spontaneous inhibitory postsynaptic currents, sIPSCs), excitability (sequential spikes), and excitatory synapse reception (spontaneous excitatory postsynaptic currents, sEPSCs) [26], indicating that external stimuli are critical for restoring NAc activity. In our study of NAc-DBS in CUMS mice, high-frequency and long-term stimuli delivery modulated the NAc D1-MSNs/dopaminergic VTA circuitry, ultimately improving the depressive-like symptoms of CUMS mice, such as increased sucrose consumption and decreased immobility time in the tail suspension and forced swimming tests. Similarly, Francis et al. reported that repeated manipulation of activity in the MSN subtypes or whole NAc can alter the depression-like responses to a social defeat stress [34]. Moreover, McCracken et al. observed resilient outcomes after repeated optogenetic high-frequency stimulation (HFS) to the whole NAc or D1-MSNs. Nevertheless, it remains unclear why HFS is effective, whereas stimulation at <50 Hz is not. The effectiveness of HFS DBS may be mediated by frequency band shifts in local field potential oscillations and mesolimbic synchrony as a result of HFS [59]. Additionally, Schmuckermair et al. [60] reported that neither single nor repeated NAc-DBS affects the depression-like behavior of WT mice, which, similar to nondepressed human subjects, do not respond to pharmacological interventions [60–62], suggesting that repeated NAc-DBS using the present stimulation conditions is only effective in deranged (i.e., pathophysiological) systems [6].

The loss of motivation and anhedonia observed in MDD can be associated with abnormalities in the reward system–dopaminergic mesolimbic pathway, in which the NAc and VTA play major roles [15]. The VTA is one of the most important anatomical substrates for drug rewards as well as natural rewards induced by food, sex, and social interactions [63]. The VTA region is a heterogeneous nucleus composed of dopaminergic and nondopaminergic neurons [64, 65]. Nondopaminergic VTA neurons are mainly GABAergic [66, 67], whereas a small percentage are glutamatergic [68]. Electrophysiological data have shown that GABAergic interneurons exert tonic inhibition over a subset of VTA dopaminergic neurons [69]. Bocklisch et al. [70] reported that NAc MSNs exert strong inhibition onto VTA-GABA neurons and functionally weaker connectivity onto DA neurons. Despite the weak direct inhibitory connection to VTA-DA neurons, D1-MSN terminal activation of the NAc could drive disinhibition of VTA-DA neurons [70]. According to reports in the literature and our data (Supplementary Figure 2), CUMS reduced action potential firing in VTA dopaminergic neurons in mice subjected to CUMS. Optogenetic phasic stimulation of VTA dopamine neurons reversed the CUMS-induced depressive-like behavior, whereas optogenetic inhibition of VTA dopaminergic neurons induced behavioral despair and decreased sucrose preference in stress-naïve mice [39, 71]. Thus, our results showed that a single trial of NAc stimuli could significantly increase GABA levels and DA neuron activity in the VTA (Figure 2). Therefore, we speculated that long-term DBS may continuously increase the excitability of D1-MSNs, which project to the VTA-GABA neurons, enhance the NAc-to-VTA connectivity, and alter the GABA neurotransmitter. Finally, the activity of VTA-DA neurons was increased substantively by modulating that of VTA-GABA neurons. Vega-Quiroga et al. also reported similar results in the regulation of VTA dopaminergic neurons via lateral septum (LS) depolarizing stimulation by K^+^, i.e., LS stimulation significantly increased dopamine and GABA, but not glutamate, VTA extracellular levels, and intra-VTA infusion of bicuculline, which is a GABA_A_ receptor antagonist, inhibited the increase in dopamine, but not of GABA VTA, levels induced by LS stimulation. By contrast, the presence of a GABA_B_ antagonist, CGP-52432, in the VTA did not produce significant changes in the VTA response to LS stimulation [72]. We obtained a similar result, i.e., Bic systemic administration blocked the antidepressant effect of NAc-DBS in CUMS mice (Figure 4). By contrast, it is also reported that acute electrical stimulation of the VTA, which is a modified version of DBS that applies short-term low-frequency programmed stimulation instead of continual stimulation, alleviates the depressive-like behavior of Flinder Sensitive Line rats, a genetic animal model of depression [73]. This may be related to the characteristics of the target nucleus stimulated by DBS.

In the central nervous system, inhibitory neurotransmission is primarily achieved through the activation of GABA receptors. Three types of GABA receptors have been identified based on their pharmacological and electrophysiological properties [74]. The predominant type, the GABA type A receptor (GABA_A_), and a recently identified GABA_C_ type form ligand-gated chloride channels, whereas metabotropic GABA_B_ receptors activate separate cation channels via G-coupled proteins. A functional GABA_A_ receptor is considered assembled from five subunits that form a pentameric chloride channel and can be inhibited by bicuculline or picrotoxin. Bicuculline is a selective and competitive GABA_A_ receptor antagonist that reduces chloride channel current by decreasing their open frequency and mean duration [24]. VTA dopaminergic and GABAergic neurons express both fast ionotropic GABA_A_ receptors (GABA_A_Rs) and slow metabotropic GABA_B_ receptors (GABA_B_Rs) [66]. The NAc→VTA input synapses onto both GABA and dopamine neurons, whereas they activate different receptors. NAc→VTA projections inhibit GABAergic neurons through GABA_A_Rs and inhibit dopaminergic neurons via GABA_B_Rs [20]. Interestingly, it has been shown that there is a segregation of the type of GABA_A_ receptors expressed in both neuronal types [42, 61, 66]. GABA_A_-α1 receptors are expressed in VTA GABAergic, but not in dopaminergic, neurons [72]. The results presented above were consistent with those of our immunohistochemistry data (Figure 6B and Supplementary Figure 4). Therefore, in our study, both the pharmacological inhibition of GABA_A_ receptor activity and genetic knockdown of GABA_A_-α1 receptor expression blocked the antidepressant effect of NAc-DBS in CUMS mice (Figures 4 and 5).

In summary, the data presented here demonstrate that long-term and high-frequency-dependent NAc-DBS can rescue the enhanced and gradual depression-related behaviors in the CUMS mouse model. The specific changes that occurred in challenge-induced neuronal activation suggest that the beneficial effects of NAc-DBS are mediated via a reward circuitry that includes the NAc and VTA. Furthermore, the enhanced GABA neurotransmitter levels and VTA-DA neuronal activity in the VTA after repeated NAc-DBS indicate that long-term alterations may be an important part of these mechanisms of NAc-DBS, leading to a substantial remedial effect of exaggerated depression-like symptoms via disinhibition of DA neurons. Although considered an invasive treatment, DBS has been proven to be safe in humans and can offer therapeutic options for these otherwise untreatable disorders. In the future, we are planning to study the molecular mechanisms of NAc-DBS in CUMS mice. At present, small-molecule compounds and their target signaling pathways have been screened in this context (data unpublished). The above-mentioned research will allow the use of drugs to replace DBS in the treatment of MDD.

## Supplementary information

**Supplementary Figure 1** Anatomical connectivity between the nucleus accumbens (NAc) and the ventral tegmental area (VTA). (A) Experimental setup: rAAV-hSyn-Cre- WRPE-PA was injected into the unilateral NAc, and rAAV-Ef1α-Dio-EYFP-WPRE-PA was injected into the ipsilateral VTA. (B) Results of NAc immunohistochemical staining. Green represents NAc neurons infected with Cre-AAV, red represents a biomarker of dopamine receptor type 1 (D1R) on NAc neurons, and blue is the nucleus (scale bar, 500 μm). (C) Results of VTA immunohistochemical staining. Green represents Cre-dependent green fluorescence (EYFP) expression in VTA neurons infected with Dio-AAV, red represents a biomarker of dopamine neurons, and blue is the nucleus (scale bar, 500 μm).

**Supplementary Figure 2** Chronic optical stimulation of VTA-DA neurons alleviates the depression-like behavior of CUMS mice. (A) Schematic representation of the oChIEF-mediated activation of VTA-DA neurons in CUMS mice. (B) Representative image showing oChIEF-tdTomato and mTH-EGFP colabeling in VTA_DA neurons. Scale bar, 50 μm. (C) Timeline for virus injection and battery of behavioral tests performed in this study. The red and blue lines represent the CUMS and chronic optical stimulation periods, respectively. (D) The sucrose preference test (SPT) was used to measure the extent of the anhedonic-like behavior during the course of the CUMS protocol and optical stimulation treatment every week (CUMS_0W: WT, 0.1122 ± 0.01724 g/g, n = 10; LED_VTA-DA, 0.09451 ± 0.01392 g/g, n = 10; *t* = 0.7971, *P* = 0.4358; CUMS_2W: WT, 0.08636 ± 0.01047 g/g, n = 9; LED_VTA-DA, 0.04901 ± 0.004766 g/g, n = 10; *t* = 3.363, *P* = 0.0037; CUMS_3W/LED_1W: WT, 0.08605 ± 0.00986 g/g, n = 7; LED_VTA-DA, 0.05914 ± 0.003845 g/g, n = 9; *t* = 2.785, *P* = 0.0146; CUMS_4W/LED_2W: WT, 0.09776 ± 0.01261 g/g, n = 7; LED_VTA-DA, 0.07096 ± 0.003433 g/g, n = 7; *t* = 2.051, *P* = 0.0628). (E) The tail suspension test (TST) was used to assess the depression-like behavior during the course of the CUMS protocol and optical stimulation treatment every week (CUMS_0W: WT, 130.1 ± 10.08 s, n = 10; LED_VTA-DA, 113.6 ± 10.08 s, n = 10; *t* = 1.155, *P* = 0.2632; CUMS_2W: WT, 130.1 ± 10.08 s, n = 10; LED_VTA-DA, 190.2 ± 8.315 s, n = 10; *t* = 4.606, *P* = 0.0002; CUMS_3W/LED_1W: WT, 130.1 ± 10.08 s, n = 10; LED_VTA-DA, 163.9 ± 16.28 s, n = 10; *t* = 1.77, *P* = 0.0937; CUMS_4W/LED_2W: WT, 156.4 ± 14.16 s, n = 7; LED_VTA-DA, 110 ± 21.72 s, n = 7; *t* = 1.791, *P* = 0.0986). (F) The open field test was used to measure locomotor activity after 2 weeks of optical stimulation (total distance: F_(2, 18)_ = 16.08, *P* < 0.0001, n = 7 animals per group; central distance: F_(2, 18)_ = 2.036, *P* = 0.1596, n = 7 animals per group; resting time: F_(2, 18)_ = 20.88, *P* < 0.0001, n = 7 animals per group; and time spent in the center: F_(2, 18)_ = 1.015, *P* = 0.3823, n = 7 animals per group). Data in D and E represent the means ± SEM; \*\**P* < 0.01, \*\*\**P* < 0.001, \*\*\*\**P* < 0.0001, *t*-test. Data in F represent the means ± SEM; \*\**P* < 0.01, \*\*\**P* < 0.001, \*\*\*\**P* < 0.0001, one-way ANOVA.

**Supplementary Figure 3** Lentivirus injection did not affect the establishment of the CUMS mouse model in each group. (A) The sucrose preference test (SPT) was used to measure anhedonic-like behavior during the course of the CUMS protocol in mice injected with i_87 into the VTA in advance. The data represent the means ± SEM; \**P* < 0.05, \*\**P* < 0.01, \*\*\**P* < 0.001, \*\*\*\**P* < 0.0001, one-way ANOVA (CUMS 0W: F_(4, 33)_ = 0.9818, *P* = 0.4309, n = 6–9 animals per group; CUMS 1W: F_(4, 34)_ = 6.566, *P* = 0.0005, n = 6–10 animals per group; CUMS 2W: F_(4, 31)_ = 14.04, *P* < 0.0001, n = 6–9 animals per group). (B) The tail suspension test (TST) was used to assess the depression-like behavior during the course of the CUMS protocol in mice injected with i_87 into the VTA in advance. The data represent the means ± SEM; \**P* < 0.05, \*\**P* < 0.01, one-way ANOVA (CUMS 0W: F_(4, 33)_ = 0.5457, *P* = 0.7034, n = 6–9 animals per group; CUMS 1W: F_(4, 33)_ = 5.502, *P* = 0.0017, n = 5–10 animals per group; CUMS 2W: F_(4, 32)_ = 6.635, *P* = 0.0005, n = 5–10 animals per group).

**Supplementary Figure 4** Confocal images of VTA slices from VTA neurons injected with LV_RNAi_87 or LV_RNAi_Vehicle. VTA_DA (Th^+^) neurons (red) were not colabeled with an anti-GABA_A_-α1 antibody (green) in mice injected with LV_RNAi_Vehicle. In turn, VTA_GABA (GABA^+^) neurons (green) were not colabeled with the anti-GABA_A_-α1 antibody (red) in mice injected with LV_RNAi_87. Scale bar, 500 μm.

## Acknowledgments

The study was supported by the Beijing Municipal Science & Technology Commission No. Z181100001518001, the Chinese Academy of Medical Sciences Initiative for Innovative Medicine No. 2021-1-I2M-034 and the Open Research Fund of the National Center for Protein Sciences at Peking University in Beijing No. KF-202104.

