## Supplementary figures and images for "NAc-DBS corrects depression-like behaviors in CUMS mouse model via disinhibition of DA neurons in the VTA"

### Supplemental Figure 1

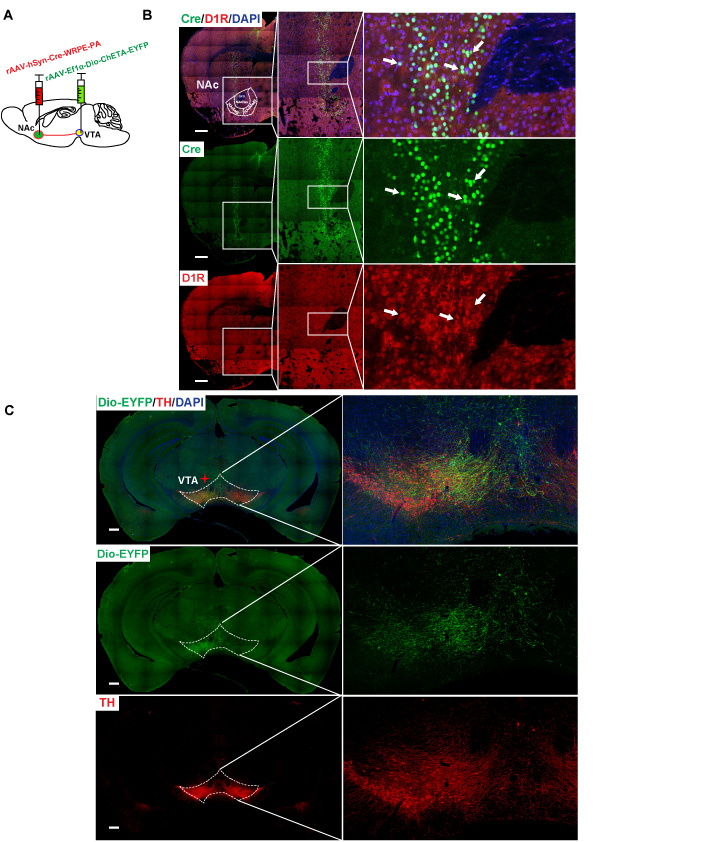

### Supplemental Figure 2

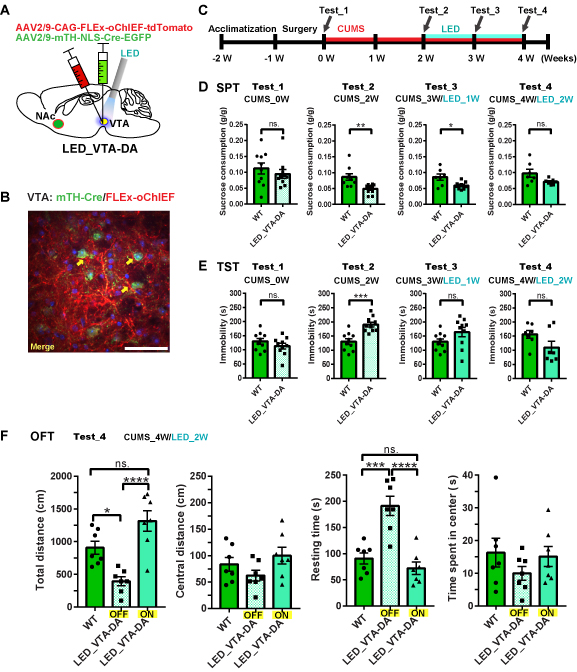

### Supplemental Figure 3

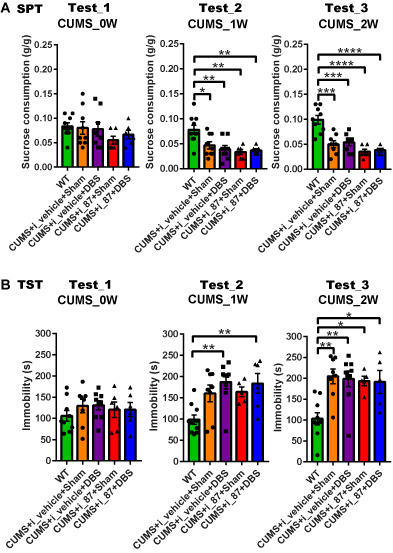

### Supplemental Figure 4

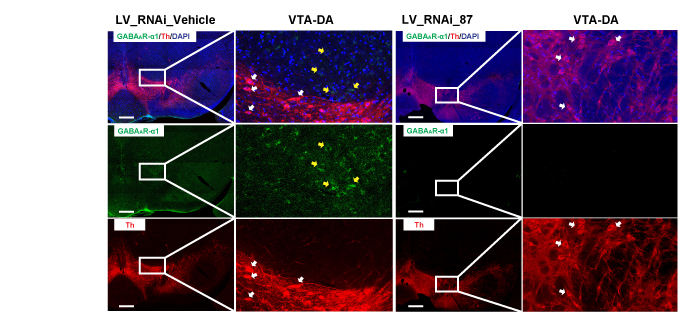
